# Genome Graphs and the Evolution of Genome Inference

**DOI:** 10.1101/101816

**Authors:** Benedict Paten, Adam M. Novak, Jordan M. Eizenga, Garrison Erik

## Abstract

The human reference genome is part of the foundation of modern human biology, and a monumental scientific achievement. However, because it excludes a great deal of common human variation, it introduces a pervasive reference bias into the field of human genomics. To reduce this bias, it makes sense to draw on representative collections of human genomes, brought together into reference cohorts. There are a number of techniques to represent and organize data gleaned from these cohorts, many using ideas implicitly or explicitly borrowed from graph based models. Here, we survey various projects underway to build and apply these graph based structures—which we collectively refer to as genome graphs—and discuss the improvements in read mapping, variant calling, and haplotype determination that genome graphs are expected to produce.

## Introduction

### The triumph of the human reference genome

The sequencing of a human genome was truly a landmark achievement (Lander et al. 2001). Over a number of years the genome assembly has steadily improved (International Human Genome Sequencing Consortium 2004; Church et al. 2011), to the point that the current Genome Reference Consortium (GRC) human genome assembly, GRCh38 (Schneider et al. 2016), is arguably the best assembled mammalian genome in existence, with just 875 remaining assembly gaps and fewer than 160 million unspecified ‘N’ nucleotides (as of GRCh38.p8).

Perhaps one reason the reference genome has been so effective as an organizing system is that the average human is remarkably similar to it. From short-read based assays, it is estimated that the average diploid human has between 4.1 and 5 million point mutations, either single nucleotide variants (SNVs), multi-nucleotide variants (MNVs), or short indels, which is only around 1 point variant every 1450 to 1200 bases of haploid sequence (Auton et al. 2015). Such an average human would also have about 20 million bases—about 0.3% of the genome—affected by around 2,100-2,500 larger structural variants (Auton et al. 2015). It should be noted that both these estimates are likely somewhat conservative as some regions of the genome are not accurately surveyed by the short read technology used. Indeed, long read sequencing demonstrates an excess of structural variation not found by earlier short read technology (Chaisson et al. 2015; Seo et al. 2016).

### Reference allele bias

Despite the relative effectiveness of the reference as a coordinate system for the majority of the genome, there is increasing concern that using the human reference as a lens to study all other human genomes introduces a pervasive reference allele bias. Reference allele bias is the tendency to under-report data whose underlying DNA does not match a reference allele (Degner et al. 2009; Brandt et al. 2015). This bias arises chiefly during the read mapping step in resequencing experiments. In order to map correctly, reads must derive from genomic sequence that is both represented in the reference and similar enough to the reference sequence to be identified as the same genomic element. When these conditions are not met, mapping errors introduce a systematic blindness to the true sequence.

In the context of genetic variant detection, this problem is most acute for structural variation. Entirely different classes of algorithm are required to discover larger structural variation simply because these alleles are not part of the reference (Sudmant et al. 2015). Furthermore, the numerous large subsequences entirely missing from the reference in turn surely contain population variation (Sudmant et al. 2015). Describing these variants and their relationships is simply not possible with the current reference model. Reference allele bias also has the potential to affect some genetic sub-populations and some regions of the genome more than others, depending on the ancestral history of the reference genome at each locus.

We believe reference allele bias is driving the field of genome inference in two directions. Firstly, with improvements in sequencing technology, the field is beginning to use unbiased de novo assembly to make inferences on individual samples. Secondly, the field is developing richer reference structures that more completely represent the variation present within the population. As a substrate for variant detection, these reference structures should mitigate reference bias by permitting a fuller complement of read mappings (see section titled ‘Genome inference with genome graphs’). These directions are not mutually exclusive; hybrid approaches will ultimately be desirable. However, this perspective focuses on reference structures, particularly for human genomics. In particular, we show how human reference-assisted variant calling is naturally progressing toward graph-based reference structures.

### Richer reference structures

The original reference human genome assembly was essentially a monoploid representation (Lander et al. 2001). The primary goal was to produce a single representative sequence albeit with regions of uncertainty—that is, a single “scaffold”—for each physical chromosome. It also included a handful of alternate scaffolds representing allelic variation, but they had no formalized relationship to the main scaffold. Recognizing that some highly polymorphic regions of the genome were particularly poorly represented by a single reference sequence, a formal model to introduce representative alternate versions of highly variable regions was added starting with GRCh37 (Church et al. 2011). Sequences in the form of kilobase to multi-megabase “alternate locus scaffolds” were described relative to the “primary” (monoploid) assembly, anchored to locations along the primary scaffolds. In the current assembly (GRCh38.p9) these cover 178 regions and total 261 sequences.

To better represent human diversity, we might imagine creating a “reference cohort”: in place of a single reference genome, a set of sequences that includes all common variation. However, representing such a cohort in the existing alternative locus scaffold system would present significant challenges. To include all alleles down to a frequency of 1%, it would require alternative locus scaffolds covering the entire primary reference genome, with hundreds of such sequences overlapping each genomic location.

While such a reference cohort is already achievable and, to some extent, derivable from public datasets like the 1000 Genomes Project (Auton et al. 2015), representing it as a collection of alternative locus scaffolds appears impractical. Firstly, the existing primary reference genome is a poor coordinate space with which to describe the other genomes. Much large structural variation is not adequately described by the coordinates provided by the primary reference. Indeed, this is already a problem with existing alternative loci scaffolds. Secondly, the alternative loci model fails to capture the fine-grained homology relationships between all the sequences (Fig. 1A). For example, when mapping a new sample into a cohort, a typical sequencing read may map equally well to many equivalent subsequences in the cohort, but this ambiguity would be illusory. Rather, this multimapping is indicative of the extensive latent structure within any nontrivial reference cohort. Each subsequence actually represents the same underlying allele. We submit that, while other models can approach it to varying degrees, this latent structure is naturally represented as a mathematical graph.

**Figure 1:**
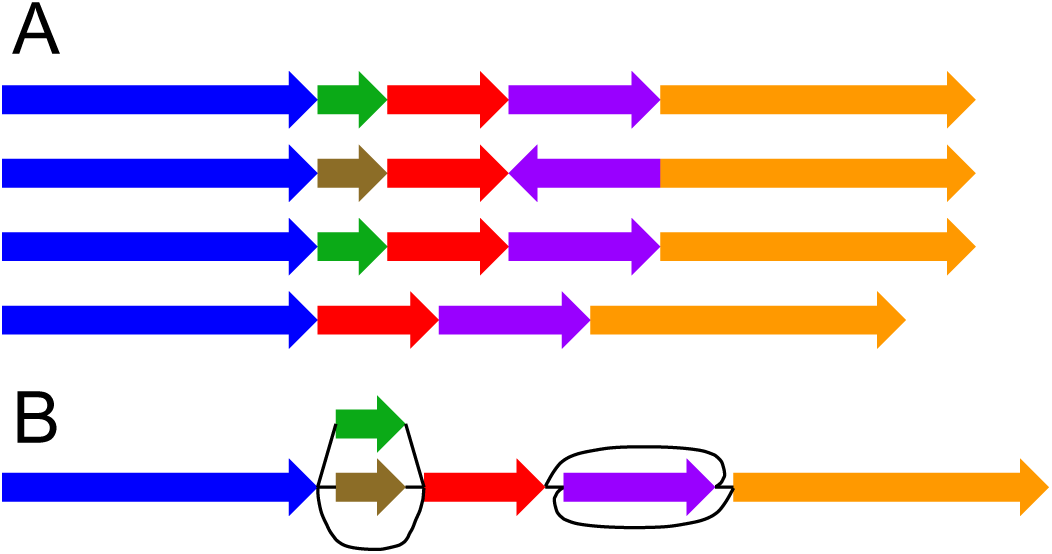
A schematic representation of two population-level reference structures. (A) A reference cohort. There is no attempt to identify homologies between the genome sequences. (B) A genome graph. Homologies are collapsed and included as alternate paths in the graph.

### Genome graphs

Graphs have a longstanding place in biological sequence analysis, where they have often been used to compactly represent an ensemble of possible sequences. As a rule, the sequences themselves are implicitly encoded as walks in the graph. This makes graphs a natural fit for representing reference cohorts, which are by their nature ensembles of related sequences (Fig. 1B).

Perhaps the simplest common graph representation is the directed graph, in which directed walks encode a nucleotide sequence. In the context of genome assembly, de Bruijn graphs (de Bruijn 1946; Pevzner et al. 2001; Zerbino and Birney 2008) a popular directed graph representation in which each node represents a *k*-mer (a unique string of length *k*) and each directed edge represents an overlap of *k –* 1 bases between the suffix of the “from” node and the prefix of the “to” node (Fig. 2A). De Bruijn graphs are a restricted class of vertex-labeled directed graphs, which are graphs whose nodes are labeled such that a directed walk can be interpreted as a DNA sequence, defined by the sequence of node labels along the walk (Fig. 2B). Alternatively, edge-labeled directed graphs are possible, in which case the nodes, rather than the edges, can be viewed as representing the intersection points between connected subsequences (Dilthey et al. 2015a).

**Figure 2:**
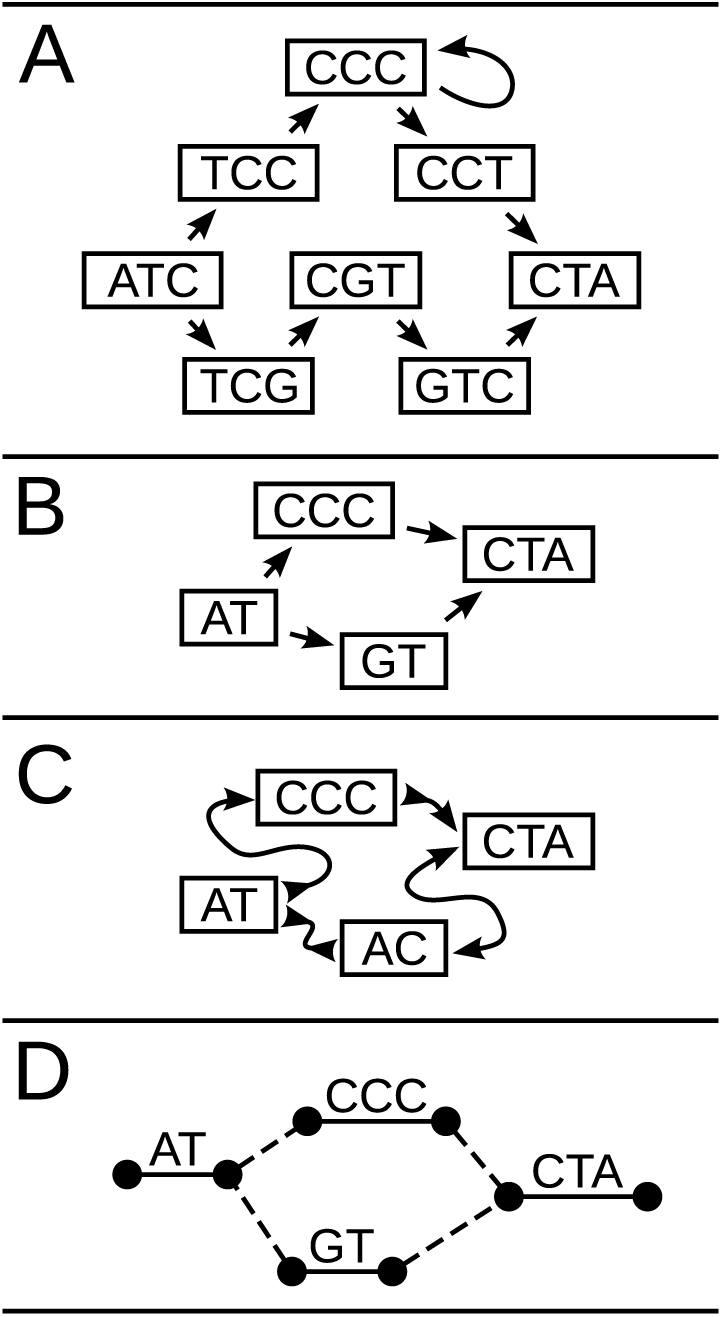
Four types of genome graphs, all constructed from the pair of sequences ATCCCCTA and `ATGTCTA`. (A) De Bruijn graph. (B) directed acyclic graph. (C) bidirected graph (aka sequence graph). (D) biedged graph (aka biedged sequence graph).

In either edge - or vertex-labeled representations, directed graphs do not fully express the concept of strand. That is, they do not distinguish the difference in reading a DNA molecule in its forward and reverse complement orientations. To express strandedness, directed graphs can be generalized to bidirected graphs (Edmonds and Johnson 1970; Medvedev and Brudno 2009), in which each edge endpoint has an independent orientation, indicating whether the forward or the reverse complement strand of the attached node is to be visited when entering the node through that endpoint of the edge (Fig. 2C). Inversions, reverse tandem duplications, and arbitrarily complex rearrangements are expressible in the bidirected representation. Such complex variation cannot be expressed in the directed graph version without creating independent forward and reverse complement nodes and storing additional information to describe this complementarity. The edge-labeled version of bidirected graphs, which we call biedged graphs (Fig. 2D), give an equivalent representation.

We call a bidirected graph in which each node is labeled with a nucleotide string a “sequence graph” (Garrison 2016; Novak et al. 2017). In a sequence graph, a DNA sequence is read out by concatenating the node oriented labels of a walk that always enters and exits each node through edge endpoints with opposite orientations. Labels are oriented such that entering through one endpoint orientation encodes the reverse complement of entering the node through the opposite endpoint orientation.

## Graphs as spatial frameworks

While sequence graphs are an effective way to compactly represent a cohort of genomes, they introduce some complications. One of these is that it is no longer trivial to define a locus on the reference. For a linear reference, it is sufficient to refer to a genetic element by its coordinate and extent along the reference sequence, but graphs admit multiple paths that may have complex relationships to each other (Paten et al. 2014). To overcome this issue we need to define a correspondence between structures in the graph and elements in the genome. In doing so, the genome graph becomes a spatial framework for organizing and comparing a population of genomes. Developing an effective spatial framework requires attending to several considerations regarding coordinates, alleles, ordering in graphs, and genome embedding, which we explore below.

### Coordinate systems

Genome graphs need a system to refer to specific positions on the sequences they contain. Ideally, such a coordinate system should convey relevant information. Computational Pan-Genomics Consortium et al. (2016) have identified desirable properties of the linear reference genome model that more general spatial frameworks should attempt to preserve. These include the notions that the genome graph coordinates of successive bases within a genome should be increasing, that coordinates should be compact and human interpretable, and that bases physically close together within a genome should have similar coordinates. On top of these properties of “monotonicity”, “readability”, and “spatiality”, Rand et al. (2016) add a further distinction between “vertical spatiality” of bases that are allelic variants of one another and “horizontal spatiality” of bases that can appear together within a single molecule. They compare two coordinate systems based on paths through a graph: one that fulfills spatiality, and another that does not fulfill spatiality but is stable under edits to the graph.

### Allelism in graphs

Many genome inference tasks involve calling alleles at sites, so genome graphs must have an operational definition of a site. Some proposals seek to derive sites from a coordinate system. One option is to fall back on the linear reference’s coordinate system and maintain a strict correspondence between positions in the graph and reference coordinates (Dilthey et al. 2015a). However, this is somewhat restrictive, possibly nullifying some of the benefits that we are seeking from genome graphs in the first place. Another proposal uses a hierarchical, recursive approach to bolt on alternative alleles to the existing reference genome system (Rand et al. 2016), which is potentially a great improvement but still has an arbitrary dependence on the existing linear coordinates.

A more graph-centered approach is to define sites based on motifs in the genome graph. In particular, it has been proposed that sites could be described with a motif called a “superbubble” (Onodera et al. 2013) (in a directed graph) or “ultrabubble” (Paten et al. 2017) (a generalization to bidirected graphs). In brief, ultrabubbles and superbubbles are directed acyclic subgraphs that connects to the rest of the graph through one source node and one sink node (Fig. 3). This motif tends to be created when new variants are added to a graph. Both superbubbles and ultrabubbles can also identify nesting and overlapping relationships involving structural variants, which makes them a more expressive definition of a site than the reference coordinate. Paten (Paten et al. 2017) shows how this site nesting is naturally described by a cactus graph, a structure that globally organizes the sites without the need for any existing reference genome. However, this approach is incomplete; not all variation is nicely partitioned into these bubbles. Moreover, it lacks the appealing simplicity of linear reference coordinates.

**Figure 3:**
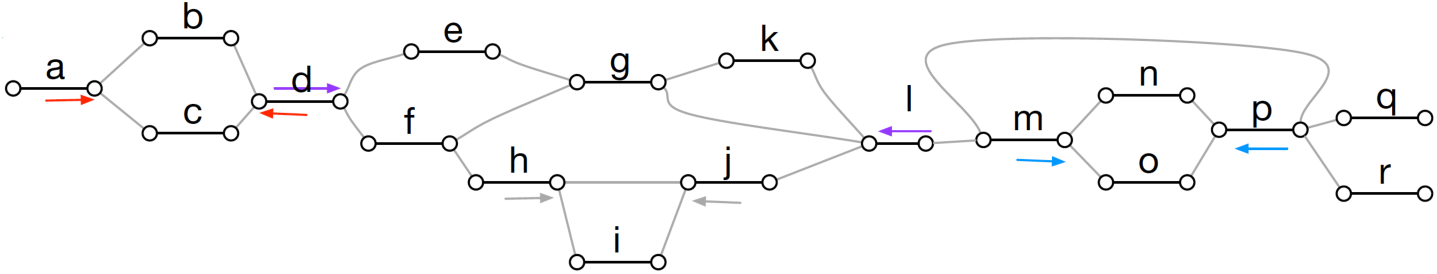
Ultrabubble sites in a biedged sequence graph. Each arrow shows the terminal node of a site. The color of the arrows indicates the node pairing. Note that the ultrabubble denoted by the gray pair of arrows is nested within the ultrabubble denoted by the purple arrows. (Reprinted from Paten et al. (2017) with permission from author.)

### Pan-genome ordering

While genome graphs do not in general provide a natural, simple coordinate system, it is possible to impose a linear coordinate system by constructing a comprehensive linear ordering of the nodes (Fig. 4) (Nguyen et al. 2015). Such a structure has been described as a pan-genome (Herbig et al. 2012; Nguyen et al. 2015). For any chromosome, such a complete ordering is potentially far more inclusive than any individual extant haplotype, in that it can include all the elements present in the population. As an actual nucleotide sequence (typically after collapsing substitutions into their major allele) this sort of linearized sequence can also potentially be closer to being a median of the genomes it represents. Nguyen et al. (2015) demonstrated useful properties of such an ordering for common bioinformatics tasks such as read mapping. They also illustrated how a linear pan-genome can be used to more inclusively visualize a set of genomes. In addition, linear orderings of genome graphs potentially have utility for creating accessible storage (ensuring good data colocalization for adjacent graph elements) (Haussler et al. 2017) and for algorithms that need to efficiently address contiguous subgraphs of a larger genome graph.

**Figure 4:**
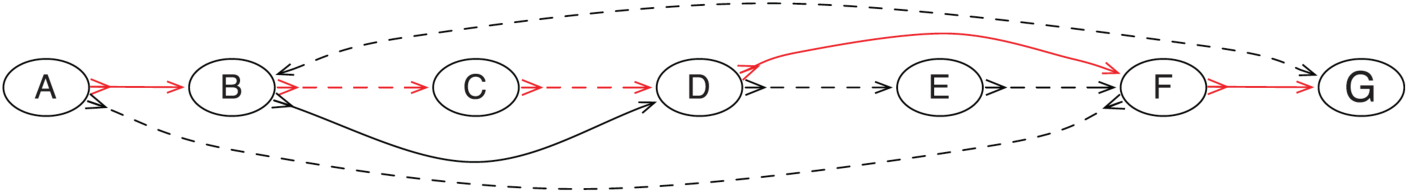
A pan-genome ordering on a graph constructed from two genomes. The red edges indicate the path of the pan-genome through the graph. The solid and dotted edges indicate the adjacencies between nodes in the two source genomes. (Adapted from Nguyen et al. (2015) with permission from author.)

### The repeatome

A significant fraction of a typical human genome is composed of highly repetitive satellite arrays (Willard and Waye 1987; Manuelidis and Wu 1978). Highly repetitive sequences are particularly prevalent in biologically important centromeric regions (Levy et al. 2007; Miga 2015), and one of the most fundamental biological structures, the ribosome, is encoded in repetitive sequence (Miga 2015; Levy et al. 2007). We need effective tools to deal with these repetitive regions—collectively, the “repeatome”—if we are to gain a full understanding of human genomics (Miga 2015).

Genome graphs can potentially allow the repeatome, not currently meaningfully accessible with short read sequencing approaches, to be analyzed in some fashion by collapsing repeats together in the graph (Paten et al. 2014). Rather than representing a repetitive region directly, the graph reference would represent the space of instances of a type of repeat across regions and individuals. In such a graph the presence of a specific repeat and its copy number can be identified by unique read mappings, but not necessarily the identity and locations of individual instances of the repeat—frequently a much more difficult problem.

Such a graph could be part of a larger graph reference, or it could serve as a special-purpose structure for repeat-specific studies. The technique could be applied to tandem repeat arrays as well as to more accessible isolated instances of repetitive elements. In the pioneering work of Miga et al. (2014), for example, probabilistic, condensed graphs for the X and Y centromeric repeat arrays were constructed, in which similar instances of a repeat were combined within the same graph node (Fig. 5). These graphs were then used to generate linear reference sequences for the two arrays (Miga et al. 2014) using a Markovian traversal, and have subsequently been used to define linear representations of the centromeres that are included in GRCh38 (Schneider et al. 2016).

**Figure 5:**
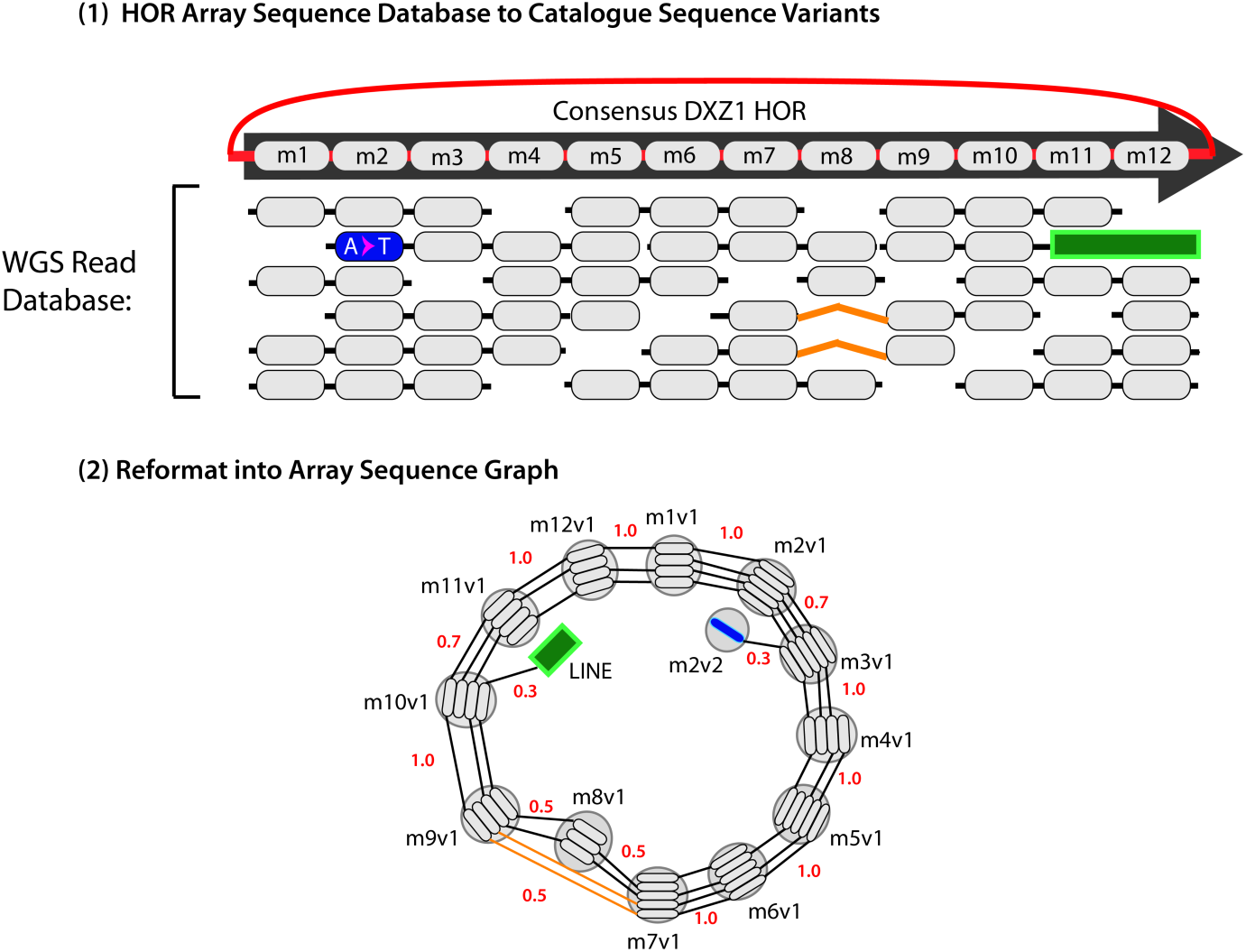
A schematic example of an “Array Sequence Graph” of the type used to construct a linearization of the DXZ1 repeat array in the X chromosome centromere (Miga et al. 2014). A collection of reads (top) shown in the context of a consensus higher-order repeat are converted into a graph representation (bottom). A cycle around the graph represents a higher-order repeat, and the individual repeat units (oblongs) are represented within each node (circles). Edges between individual repeat units represent phasing information from input reads. Transitions between nodes are annotated with probabilities. (Adapted from Miga et al. (2014) with permission from author.)

### Hierarchy

Extending the idea of collapsing repetitive sequences, the work of Paten et al. (2014) showed that it is not necessary to pick a single genome graph to act as a reference. Rather they argued that a hierarchy of graphs related by graph homomorphisms (a projection function from the nodes of one graph to the nodes of a more collapsed representation of that graph) could be constructed in which progressively more collapsed versions of the same underlying set of genomes could be constructed and related (Figure 6). In the most collapsed graph in the hierarchy repetitive sequences could be fully collapsed, while the least collapsed graph in the hierarchy might represent the input set of haplotypes as disjoint sequences. Intermediates in this hierarchy could represent a more typical monoallelic representation of the genomes. The constructed hierarchy would have the property that mapping a subsequence to an element in a graph in the hierarchy automatically implies the mapping of the subsequence to all the more collapsed versions of that element in the more collapsed graphs in the hierarchy. In this way, mapping a sequence to a specific repeat instance would also identify the sequence as mapping to the canonical copy, classifying it as an instance of the repeat type.

**Figure 6:**
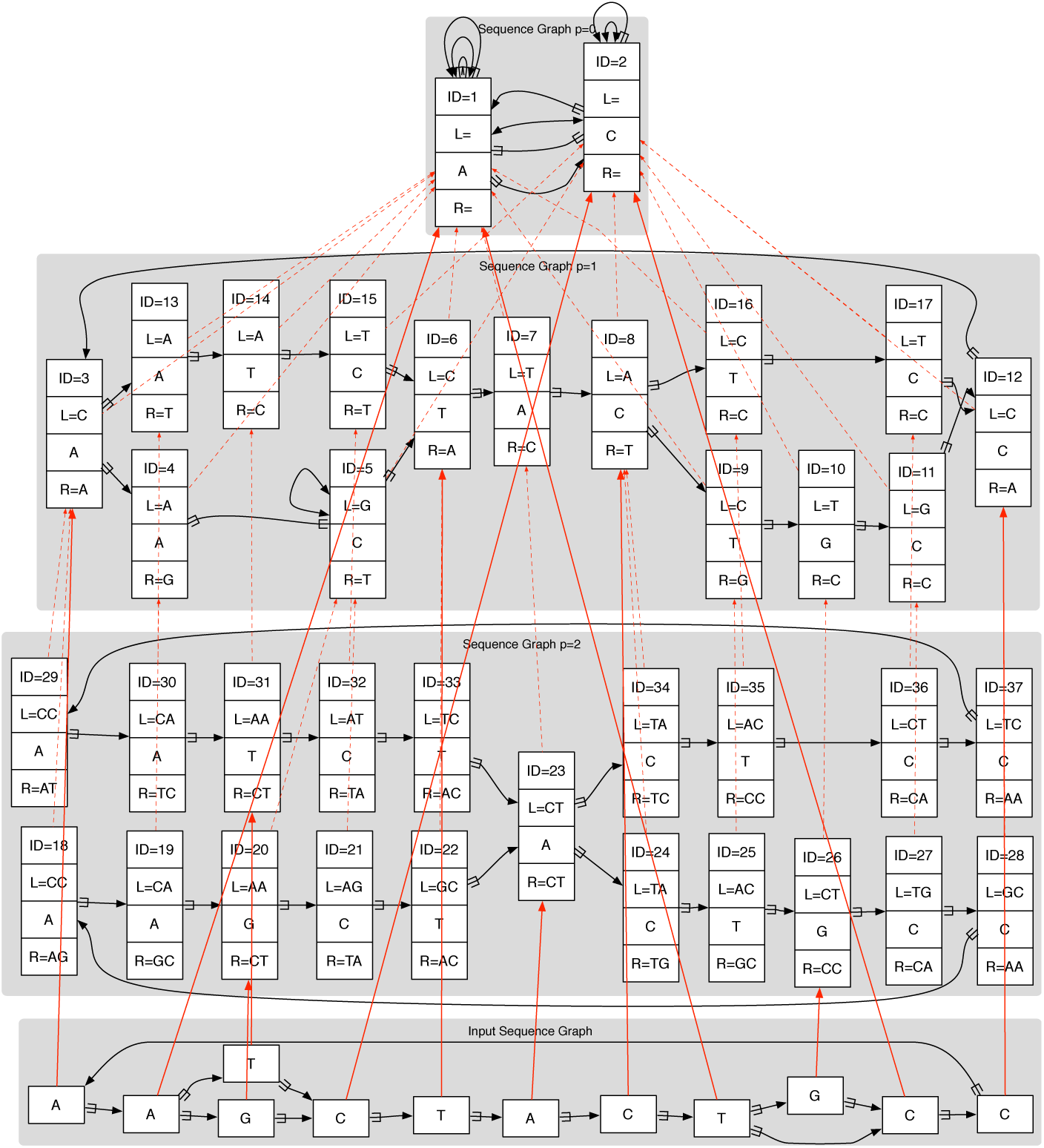
A reference genome graph hierarchy (most collapsed graph at the top, less collapsed lower down) with an input graph (bottom) mapped to it. All the graphs in the reference hierarchy are de Bruijn graphs. Dotted red lines show projections between graphs in the hierarchy, while solid red lines show mapping of the input sequence graph into the hierarchy. Here each node has a unique ID and the L and R strings represent flanking contexts mapping strings required for unique identification. (Reprinted from Paten et al. (2014) with permission from author.)

### Haplotype embedding

The number of paths through a genome graph increases combinatorially with the number of alternate alleles it includes. However, many of these sites are in tight linkage disequilibrium, so the number of paths that actually occur in the population increases much more slowly (Fig. 7). It can be useful for read mapping and variant calling to distinguish between paths have actually been observed and others, which are likely rare or absent in the population. One solution is to store allele frequency and linkage information in the genome graph by embedding a population cohort as a set of walks through the graph, each of which represents a haplotype. Such a sequence graph with embedded walks is known as a *variation graph.* This is also an instance of the graph hierarchies described above in which there is homomorphism from the population cohort onto a constructed genome graph.

**Figure 7:**
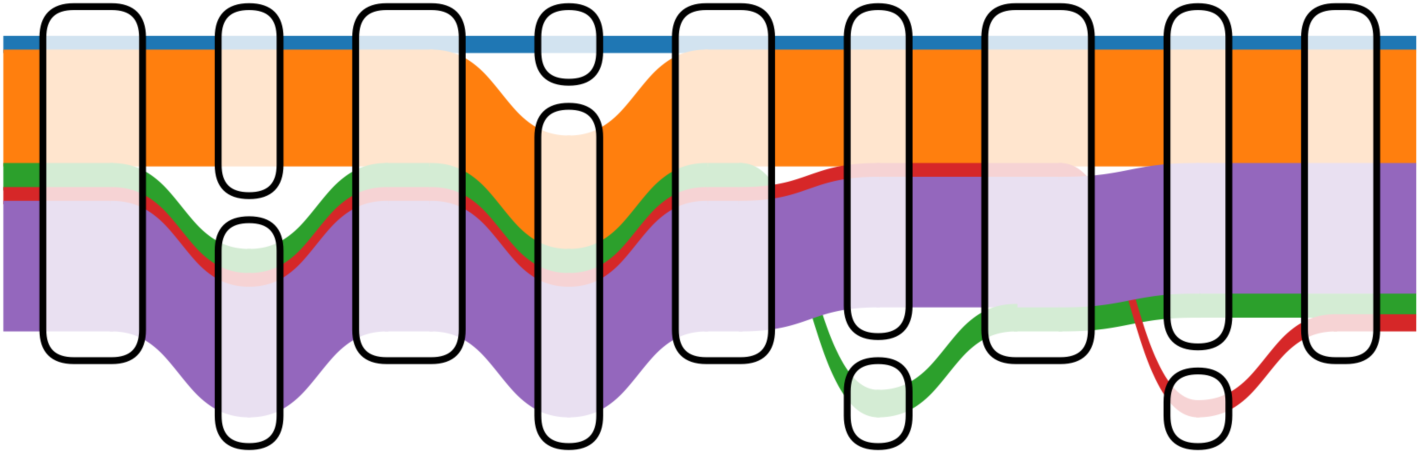
Distinct 1000 Genomes Project haplotypes embedded within a variation subgraph. Haplotypes are shown as colored ribbons with width proportional to the log of their frequency. The number of possible paths traversing left-to-right is 16, but only 5 are observed in 1000 Genomes because of linkage disequilibrium. Figure based on prototype by Wolfgang Beyer (personal communication).

A major challenge to implementing embedded haplotypes is making the data structure for storing the cohort compact enough to fit in memory while also keeping the data available for computation. In linear references, the Positional Burrows Wheeler Transform (PBWT) was developed in response to this challenge. The PBWT is a transform of a matrix of binary haplotypes that is highly compressible and supports efficient haplotype search queries even when compressed (Durbin 2014). Building on this idea, the recently developed graph-PBWT (gPBWT) is a similar succinct data structure that supports simultaneous compression and efficient haplotype queries on haplotypes embedded as walks within a variation graph (Novak et al. 2016). It appears practical to store entire population cohorts for thousands of genomes in such an auxiliary structure, which is already implemented within the xg software (https://github.com/vgteam/xg).

## Read mapping in genome graphs

One of the most important use cases for a genomic reference is as a target for read mapping.

Read mapping tools usually rely on one of two indexing approaches, in order to quickly find the best mapping locations for a read in a reference. Some tools, like MOSAIK, use *k*-mer-based indexing, in which short *k*-mer sequences from the reference are related to their locations using a hash table or other traditional key-value data structure (Lee et al. 2014). Other tools, such as BWA-MEM, use Burrows-Wheeler Transform-based approaches, in which the reference is stored in a succinct self-indexed data structure optimized for substring search (Li 2013). However, these approaches have traditionally been implemented against monoploid, linear reference genomes, resulting in a mapping task where each read is placed at one or more linear coordinates. Mapping against more complex, nonlinearized pangenomes presents unique challenges.

Several notable read mapping tools (or more integrated tools that internally map or process reads) are described in Table 2. Both linear-reference-based and graph-based tools are represented.

**Table 1:**
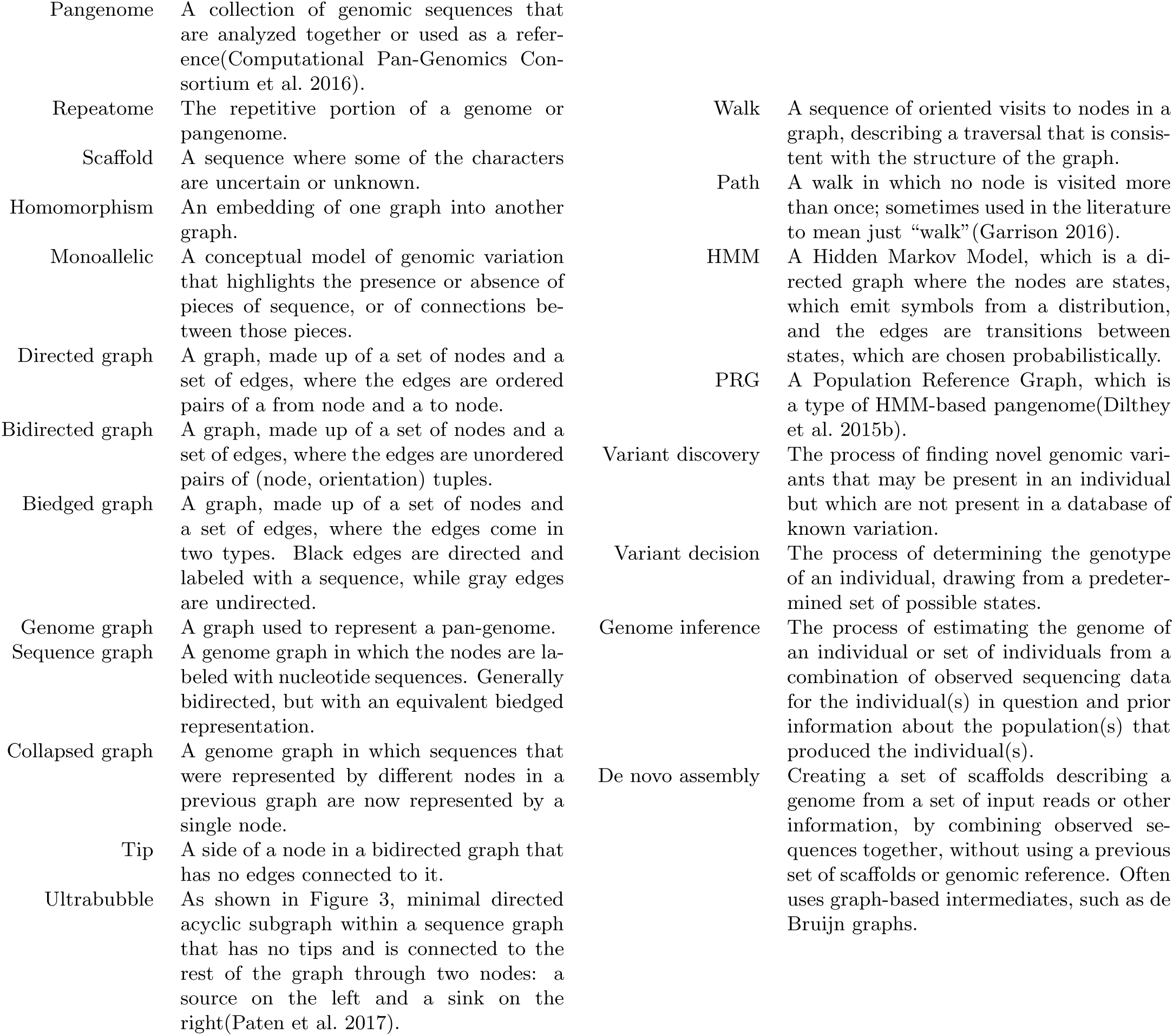
A glossary of terms used in this perspective.

**Table 2:**
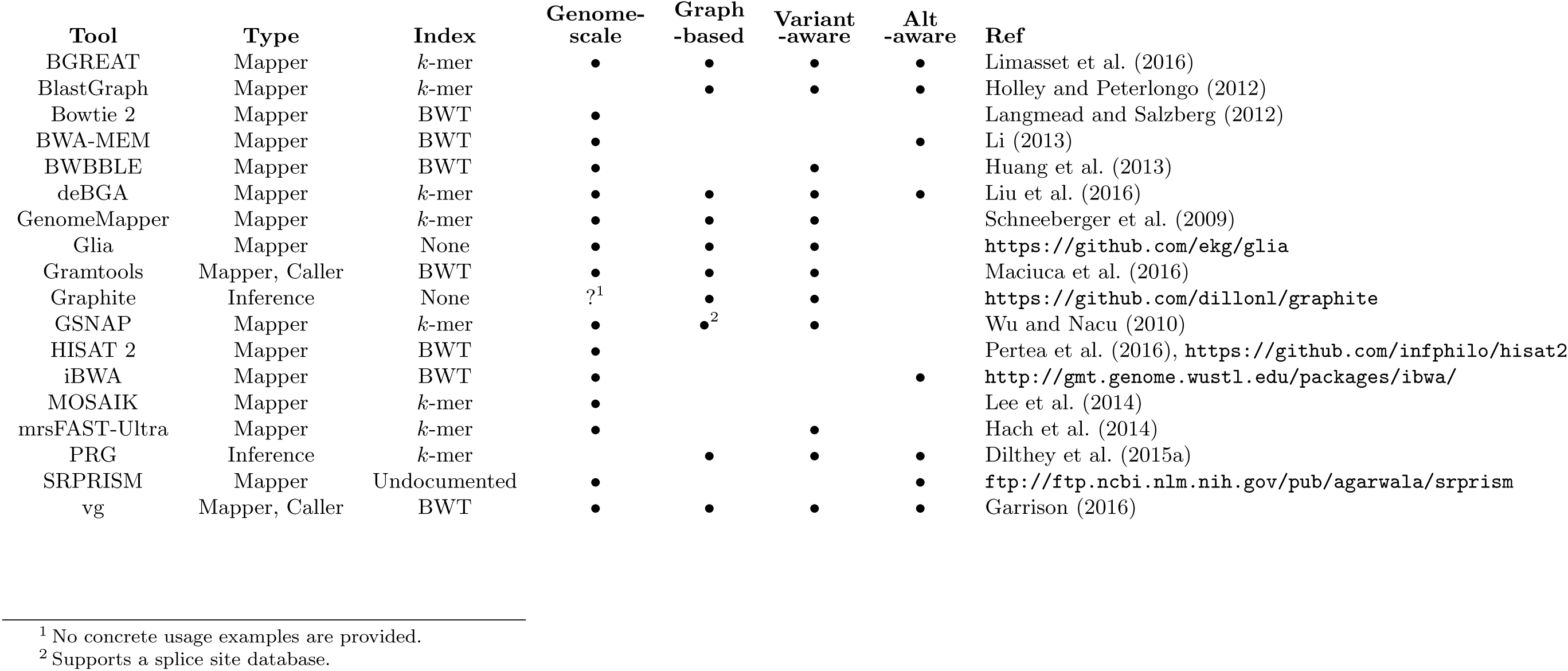
A comparison of various read mapping and processing tools. Tools come in several types. “Mapper” tools perform read mapping; “Caller” tools also perform variant calling; “Inference” tools take in reads and perform specialized or integrated genome inference tasks. The “Scale” column denotes the size of reference that each tool has been demonstrated on; some tools have been demonstrated or are marketed as being suitable for genome-scale analyses, while others are intended for regions such as the MHC or do not give any concrete usage examples. Tools are also graded on the presence or absence of several traits. “Genome-scale” tools have been demonstrated on or are marketed as suitable for the analysis of entire human genomes. “Graph-based” tools make use of an internal graph representation of variation or support adjacency information. “Variant-aware” tools are capable of accounting for point or small-scale variation in a reference. “Alt-aware” tools are capable of accounting for larger-scale replacements or for the presence of complete secondary contigs in a reference. Note that, among the tools described here, graph-based tools are always variant-aware, and only graph-based tools are both variant-aware and alt-aware.

### Alt-aware mapping

The current human genome assembly, GRCh38 (Schneider et al. 2016), contains alternative loci which provide different versions of sequences already represented in the primary assembly (see the section “Richer reference structures”). Mappers which can make sense of these additional representations, either as special linear sequences or as components of a graph, are marked as “alt-aware” in Table 2.

While alt-awareness naturally falls out of many graph-based pangenome representations, it is also possible to achieve in a linear framework. BWA-MEM is one example of a linear-reference-based, alt-aware mapping tool (Li 2013; Church et al. 2015). Although not formally described in the literature, the alt-aware feature of BWA-MEM uses a two-step process (https://github.com/l" h3/bwa/blob/1f99921b73237203e5772bee5a8c7a254c6bcbce/README-alt.md. In the first step, reads are mapped to the primary and alt sequences, as normal, but with with special rules for mapping quality calculation and primary/ secondary/supplementary status assignment. In the second step, alignments between alt and primary sequences are employed to project alignments into the primary sequence space, and to update mapping qualities in light of how the primary and alt sequences fit together. This approach produces alignments to GRCh38, in the traditionally-used primary sequence coordinate space, but making some use of the alt loci, and there is some preliminary evidence to suggest that the resulting alignments may be better than those obtained using BWA-MEM on only the primary sequences.

If one of a person’s two haplotypes in a region is much closer to an alt sequence than the primary sequence, the projection onto the primary sequence’s coordinate space will at best lose information and at worst interfere with variant calling. One feature of BWA-MEM’s alt-aware mode is that it also outputs alignments in the coordinate spaces of the alts, to allow variant calling in alt coordinates. However, how to use these alt-coordinate-space alignments to produce a combined set of variant calls across the linear coordinate spaces most appropriate for an individual is still an open question (but see Dilthey et al. (2015a)).

### Extending mapping to genome graphs

Some graph-based approaches, like Gramtools (Maciuca et al. 2016), admit only certain highly structured graphs. This allows them to be “variant-aware”—to account for small-scale variants when computing read alignments—with a controlled amount of additional complexity over linear-reference-based approaches. Other approaches can handle more general directed acyclic graphs, or even bidirected, cyclic sequence graphs. Extending local alignment, indexing, and distance measurement to graphs—and especially to complex graphs—has proven to be a challenge, and the tools in Table 2 have approached these problems in various ways.

### Local Alignment

While extending fully-ordered dynamic programming algorithms to partially-ordered directed acyclic graphs is relatively simple (Lee et al. 2002), sequence graphs are not restricted to directed acyclic structures. Some biological structures, such as inversions, duplications, or highly variable copy number, might most naturally be represented using the bidirectedness of the sequence graph formulation, or by adding cycles to the reference. In vg, one of the tools that supports these more complex structures, cyclic bidirected graphs are “unrolled” and “unfolded” to make directed acyclic graphs that are amenable to partial-order alignment, with the results transformed back into the original graph’s coordinates (Fig. 8) (Garrison 2016). In deBGA, another such tool, complex structures in the de Bruijn graph index are flattened out when alignments are articulated against individual linear haploid assemblies (Liu et al. 2016).

**Figure 8:**
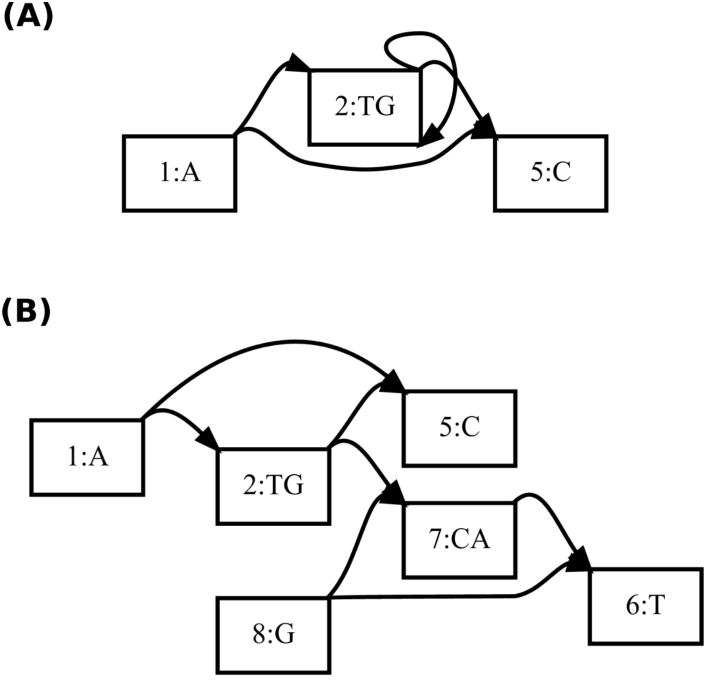
A bidirected sequence graph (A) being unfolded into a directed acyclic graph (B), in preparation for partial-order alignment. Node 6 is a reversed view of node 1, node 7 is a reversed view of node 2, and node 8 is a reversed view of node 5. (Reprinted from Garrison (2016) with permission from author.)

### Indexing

When working at the scale of whole genomes, the problem of extending indexing strategies to graphs becomes very important. Read mapping tools need to narrow down the reference genome when mapping a read, usually to one or a few regions containing candidate hits. As with tools for mapping to linear genomes, sequence graph mapping tools can be divided into those that use succinct self-index approaches and those that use *k*-mer lookup table approaches. On the succinct self-index side, one notable example is Gramtools’s vBWT, where the graph itself is represented as a modified BWT (Maciuca et al. 2016). This approach is quite elegant, but it limits the structure of the graph to one that merely represents successive sets of alternatives. On the other end of the spectrum, vg, which uses the GCSA2 graph indexing library, is able to represent arbitrary sequence graphs in its index, although index size grows combinatorially with local graph complexity (Garrison 2016; Sirén 2017). One interesting approach to controlling index size is illustrated in HISAT 2 (Pertea et al. 2016); although its approach is not yet described in the literature, the tool is built around a collection of small graph self-indexes, rather than a single large index.

One of the most interesting graph-based tools using a *k*-mer-based index is the “population reference graph” (PRG) method of Dilthey et al. (2015a). This tool uses a Hidden Markov Model (HMM), with emission distributions over sets of *k*-mers. The *k*-mers found in the reads are used to infer a pair of representative linear haploid genomes for a sample, denoted as paths through the HMM. These sequences are then used for re-alignment with the succinct self-index-based BWA-backtrack, in order to produce a final set of read alignments. More broadly, de-Bruijn-graph-based tools, such as deBGA, are marked as using *k*-mer-based indexes, because the nodes in a de Bruijn graph are identified and looked up by *k*-mers (Liu et al. 2016).

### Distance Measurement

Finally, there is also the question of distance measurement in graphs, for the purposes of paired end resolution. A serious contender in the read aligner space must deal with paired end reads in a reasonable way, which is challenging in a genome graph because calculating the distances between mappings, or even the relative orientations of mappings, is no longer trivial.

There are two major approaches described in the literature: doing paired-end resolution in the space of some (potentially inferred) linear sequence, and doing paired-end resolution using a graph-based distance metric. The PRG system and deBGA do paired-end resolution in the space of individual sequences: the generated pair of sequences used for BWA realignment in PRG, and the linear reference sequences embedded in the de Bruijn graph in deBGA (Dilthey et al. 2015a; Liu et al. 2016). A graph distance metric is used for paired end resolution in vg, which can serve as an example of that approach, although the implementation does not currently consider the relative orientations of paired reads (Garrison 2016). Some projects, like HISAT 2, have not yet documented how their paired-end distance calculations are performed in a graph, while other projects, like Gramtools, do not yet implement paired-end alignment (Maciuca et al. 2016). Overall, paired-end resolution in graph reference structures is a relatively open problem.

## Context Mapping

In addition to graph-based mapping methods based on traditional dynamic-programming alignment, there has also been interest in alternative notions of mapping, building on the notion that a position in a sequence graph reference can have different semantics than a position in a linear reference. Precedent for this idea is found in the notion of flanking sequences for SNPs in dbSNP (Sherry et al. 2001). Originating before the first release of the human genome assembly, dbSNP was not designed around a linear reference sequence. Rather than requiring variation to be submitted by coordinate in a reference genome, variation originally instead had to be submitted along with flanking sequence, describing the context in which the variation was observed by the submitting laboratory (Sherry et al. 2001). These observed flanking sequences (perhaps obtained by Sanger sequencing), rather than a VCF position, defined the variant (Sherry et al. 2001).

In a sequence graph, the sort of offset-based coordinates that are commonly used in linear references can become unwieldy. It may be convenient to instead think of graph position as not defined by a coordinate but rather by a context. Instead of finding the coordinates at which a read maps and then considering a string-to-string alignment, one can assign each position in a read to a position in a sequence graph, based on correspondences between their contexts. One particular formulation of this idea, “context schemes”, defines a mathematical formalism of non-redundant context assignment that ensures unambiguous mapping of individual positions, and demonstrates a heuristic, potentially practicable algorithm on linear references (Novak et al. 2015). Work dealing with nontrivial graphs includes the “fuzzy context-based search” approach of Leonardsen (2016) and the small sequence graph reference structure examples of Paten et al. (2014) (see Figure 6 for an example). It is anticipated that further research in this area could yield useful practical implementations, because the correspondence between context and position identity in a graph is quite natural. However, sensitivity challenges remain: concepts of spatial closeness, even between adjacent positions, are difficult to translate into a context-based framework, so reads that must be placed by integrating information across the read sequence present a particular challenge.

## Genome inference with genome graphs

The primary advantage of resequencing with a reference genome as opposed to de novo assembly is that it greatly simplifies the process of genome inference. Where assembly needs to discover the entire genomic sequence, reference-based resequencing only needs to discover a sample’s differences from the reference. Intuitively, genome graphs should provide even further advantages of the same kind. The graph contains not only the sample’s approximate sequence, but also many of its specific variants. This simplifies much of the genome inference process from discovery to decision.

Despite these theoretical advantages, research on variant calling using genome graphs is still relatively nascent. In contrast, many successful methodologies have been published for calling variants using the linear reference genome (Nielsen et al. 2011). In order to be useful in practice, genome graphs must be able to translate their promised reduction in reference bias into measurable improvements in variant calling over established methodologies. Accordingly, developing variant calling algorithms for genome graphs is an important research frontier.

The leading linear reference-based variant calling tools in use today are all based on probabilistic models of sequencing data (Nielsen et al. 2011). This approach has several advantages. Modern sequencing technologies all attempt to quantify the uncertainty in their base calls. Probability models provide a natural framework to incorporate this uncertainty into genotype calls, and they allow algorithms to estimate uncertainty about genotype calls for downstream analyses.

The actual probability models used in these tools vary, but they share a common Bayesian structure. For a genotype *G* and a set of read data *D,* the posterior likelihood of a genotype call is *P*(*G*|*D*) ∝ *P*(*D*|*G*)*P*(*G*). The first factor, *P*(*D*|*G*), is based on a generative model of the read data. The second factor, *P*(*G*), is the prior probability of a genotype, which can be flat or based on population information.

### Genotype likelihoods in genome graphs

In genome graphs, genotyping consists of two distinct tasks: decision between the variants present in the graph structure a priori and discovery of novel variation. These tasks require different inference processes, so it is not surprising that every technique we are aware of uses a different strategy for each.

Of these tasks, the genotype decision process is more novel to genome graphs. Recall that genome graphs are constructed so that a haplotype can be represented as a walk through the graph, modulo a few rare or private variants that are in the haplotype but not in the graph. Thus, for a diploid sample we can essentially frame the genotype decision process as choosing two walks through the graph that contain the sample’s variants, possibly phasing arbitrarily in the process (Fig. 7). This naturally leads to a path-based methodology for variant calling.

Several path-based variant calling methods have already been developed. PRG uses an HMM to compute the maximum likelihood pair of paths through its graph (Dilthey et al. 2015a). BayesTyper generates local paths through nearby variants. In both cases, the genotype likelihood is based on the *k*-mer content of the reads. Both *k*-mer methods forgo read mapping, which makes them computationally efficient and avoids some conceptual ambiguities with aligning to graphs. However, *k*-mer models are also inherently susceptible to sequencing errors. The PRG group has also developed a specialized genotyping algorithm for the HLA locus that is based on read alignments, called HLA^*^PRG (Dilthey et al. 2016). This algorithm has high accuracy, but its computational demands are too high to scale the approach genome-wide. Finally, the vg suite includes a nascent variant calling tool that is primarily site-based instead of path-based. It defines sites with ultrabubbles, and then defines the alleles as the set of paths through the ultrabubble. vg calls variants from read alignments with a count-based heuristic, which takes advantage of vg’s flexible aligning capabilities but currently lacks a sophisticated error model.

Compared to genotype decision, the variant discovery process in graphs more closely resembles traditional genome inference. In fact, PRG and BayesTyper both use existing variant calling tools to discover variants. BayesTyper adds the discovered variants to the graph as candidates for the path-based inference, whereas PRG converts its maximum likelihood paths into candidate linear reference sequences upon which to discover novel variants; vg uses read pileups from its alignments to augment the graph so it can then use the same model for genotype discovery as decision.

In all cases, these tools sidestep some of the difficulties in generalizing the read mapping-based models that have been successful with linear reference genomes. PRG and BayesTyper both avoid mapping reads to a graph reference by using *k*-mers. HLA^*^PRG restricts is focus to a small section of the HLA locus. The current version of vg uses heuristics in place of a true probability model for its mapped reads. We expect that true generalizations of traditional models will be a significant advancement for the field, particularly for segregating structural variants. The fact that sequence graphs make it possible to align and model known structural variants in the same manner as other types of variation may obviate the need for separate algorithms from SNV callers (Medvedev et al. 2009).

### Haplotype priors on genome graphs

As graph-based genome inference matures, it will be necessary to develop effective population priors. This need is perhaps more urgent with graphs than with linear references, because, as described above (see Haplotype Embedding) graphs can admit many paths that are not biologically meaningful. However, the novel path-based methodologies described above also represent a clear opportunity. Since the paths already combine information across sites, they provide a natural setting for incorporating linkage disequilibrium information into the prior, which is then a prior over haplotypes rather than genotypes. Doing so could mitigate the effect of the non-biological paths and perhaps even improve the accuracy of genome inference. Basic research in this vein is already emerging. A graph generalization of the Li and Stevens population model using the gPBWT described above has been developed (Rosen et al. 2017). These or similar techniques could provide the infrastructure for efficient computation of population haplotype priors.

## Future Opportunities and Challenges

We foresee reference cohorts replacing linear references, and that graphical models of reference cohorts will replace linear sequence assemblies as the space in which genomics is done. In the short term, some of the benefits of a more comprehensive reference cohort can be gained by resequencing against the full GRCh38 assembly, using software like BWA-MEM that properly handles alternative loci. We believe that extending existing software to fully support the alternative locus sequences of GRCh38 will lay the groundwork for full sequence graph support in the future.

Over the long term, we anticipate the development of an official reference cohort and graph based reference that embeds this cohort. We imagine regular releases that progressively add to this structure, and that this will be overseen by an independent group, like the Genome Reference Consortium. Such an official graph could provide a truly universal coordinate system encompassing not only a single linear assembly but also the global stock of variant information from projects like 1000 Genomes. While there are benefits to having tailored approaches for specific applications, we believe that, on balance, a greater degree of standardization would be a boon in this area. Currently, too much effort is duplicated in having each group build their own partial graph-based compendia of human variation. We believe strongly in the benefits of having a concrete, comprehensive genomic data structure upon which to base our discourse. Accordingly, there is a need to generate some consensus about the best method for constructing genome graph references.

Moreover, we expect and hope for community convergence on a universally-applicable graph formalism and exchange format. We anticipate that fully bidirected sequence graphs will be the ultimate winner, because of their ability to represent inversions, duplications, and translocations in a way that is natural and that accounts for gene flow in and out of such structures (Navarro et al. 1997). However, we anticipate that, in an official human sequence graph reference, these more general features would be used sparingly, for describing large-scale rearrangement events, because it is often convenient to work with graphs that are locally directed and acyclic. Moreover, a large fraction of variation is amenable to a directed acyclic representation. Standardizing on an exchange format for graphs, meanwhile, would allow hard-won insights about how to store, retrieve, and process graph data at species scale to make their way into new software from the reference interface inwards.

Perhaps the biggest challenge in migration to genome graph references beyond the development of the necessary data science and technology is the inertia that has developed around the current human reference assembly. As such a linchpin data structure a tremendous amount of tooling and data relies upon the current reference. We advocate the development of production quality toolkits for genome graphs, that, much like HTSlib and SAMTools (Li et al. 2009) for working with the current reference, make it easy to work with genome graph technology. We hope vg will become such a toolkit.

Toolkits will facilitate the transition, and, coupled with a progressive, incremental approach, make adoption significantly more likely. We recognize that, just as previous reference genome assemblies tend to hang around, so will the current reference human genome assembly even in a future where a significant fraction of work has switched over to using reference genome graphs. We anticipate the need to maintain translation between linear reference assemblies and genome graphs, and in particular the need for tools to lift over annotations between existing assemblies and genome graphs, just as we do today between updated assembly releases.

In the area of genome resequencing, the trend is, as always, towards longer read sizes (Jain et al. 2015), cheaper sequencing (Jain 2015), and larger read separations (Putnam et al. 2016; Coombe et al. 2016). We anticipate that these advances will allow de novo assembly to become a more practical resequencing analysis approach, and will, overall, increase the accessibility of large-scale structural variants. Although the importance of a reference for ordering and orienting short reads may wane as long read technology improves, the importance of a reference compendium of structural variation will increase as such variation becomes more detectable. Furthermore, the importance of a reference that provides information on known variable bases will increase as higher-read-error-rate sequencing methods become more widely used.

In the area of read alignment, we recommend more research into alternative problem formulations, rather than just straightforward extensions of read mapping to graphs. Instead of finding a single best alignment of a read to a graph, for example, it would be useful to be able to obtain a collection or distribution of such alignments, which would account for different possible paths in the graph from which the read may have been generated. When there are multiple decision points in an alignment, and especially when reads are long, a single “mapping quality” value may no longer be a sufficient description of alignment confidence. Carrying more detailed uncertainty information about read mapping through to the variant calling or genome inference step of a resequencing pipeline should improve overall performance. It would also be useful to explore the idea of context-based mapping more thoroughly; we anticipate that effort spent developing more practical and efficient implementations of the basic idea would not be wasted.

Future genome inference methodologies must capitalize on these developments and translate them into improved accuracy. We have already suggested some directions for this research. Inference models that have already proven successful on linear genomes should be generalized to graphs, and future models should take advantage of new path-based variant calling methodologies to incorporate linkage disequilibrium into the population prior. Ultimately this will result in the combining of phase imputation and variant inference into one model and optimization process, allowing sampling from the posterior of genomes rather than simply genotypes. When these techniques mature, we expect that they will outperform existing genome inference tools, especially in cases involving structural variation and high polymorphism. Moreover, the transition from discovery to decision processes will require less from the data to infer common variants. Thus, these methods will also serve the economic imperative of maintaining accuracy while reducing coverage requirements. This will in turn enable projects at larger scales. Conversely, we believe that there is more useful information to be extracted from existing sequencing samples, and that existing short read technologies may yet be used to generate significantly more comprehensive genome inference when compared against a comprehensive genome graph. In making a cost-benefit analysis of sequencing technologies, we believe the development of reference genome graphs and large population cohorts, essentially as a better prior for inference, may significantly alter the calculation.

In this perspective we have focused on genome graphs and their likely impact on genome inference and genetic discourse. However it is important to recognize that genome graphs could have a similarly dramatic impact on transcriptomics and epigenetics. While beyond the scope of this discussion, the natural extension of the reference genome to incorporate a more complete map of variation opens the possibility of directly linking transcript expression and epigenetic marks with specific alleles, so creating an integrated structure for relating these other omics types to underlying genomic variation.

The developments described in this perspective indicate a rapidly growing ecosystem of tools and methods for genome inference using graph-based reference structures. As a whole, graph-based genome inference promises to mitigate the problem of pervasive reference bias, through effective incorporation of reference cohort information. This will have an impact throughout genomics, as previously intractable forms of genetic variation become assayable with the efficiency of routine resequencing experiments. For some time, researchers and ethicists have warned against the health care disparities that basing genomic studies on European populations could cause (Need and Goldstein 2009). Now evidence is accumulating that such disparities are already occurring in personalized medicine (Petrovski and Goldstein 2016). Reducing reference bias is an important step toward remedying this problem, and graph genomes are the most promising proposal to do so.

## Acknowledgments

We thank Karen Miga and Wolfgang Beyer for providing help and figures. We thank David Haussler, Richard Durbin, Gil McVean and members of the reference-genomes task team of the Global Alliance for Genomics and Health for many useful discussions. We thank Daniel Zerbino and the two other anonymous reviewers for their insightful comments and feedback. We thank Glenn Hickey, Charles Markello, Yohei Rosen, Eric Dawson, Mike Lin and all contributors to the vg software package, which is driving much of our work forward. This work was supported by the National Human Genome Research Institute of the National Institutes of Health under Award Number 5U54HG007990 and grants from the W.M. Keck foundation and the Simons Foundation. The content is solely the responsibility of the authors and does not necessarily represent the official views of the National Institutes of Health.

## Disclosure Declaration

The authors of this manuscript include authors of the vg Garrison (2016) and Glia tools described above. The authors declare no other conflicts of interest.

